# Improved integration of single cell transcriptome data demonstrated on heart failure in mice and men

**DOI:** 10.1101/2023.03.15.532742

**Authors:** Mariano Ruz Jurado, Lukas S. Tombor, Mani Arsalan, Tomas Holubec, Fabian Emrich, Thomas Walther, Andreas M. Zeiher, Marcel H. Schulz, Stefanie Dimmeler, David John

## Abstract

Biomedical research frequently uses murine models to study disease mechanisms. However, the translation of these findings to human disease remains a significant challenge. In order to improve the comparability of mouse and human data, we present a cross-species integration pipeline for single-cell transcriptomic assays.

The pipeline merges expression matrices and assigns clear orthologous relationships. Starting from Ensembl ortholog assignments, we allocated 82% of mouse genes to unique orthologs by using additional publicly available resources such as Uniprot, and NCBI databases. For genes with multiple matches, we employed the Needleman-Wunsch global alignment based on either amino acid or nucleotide sequence to identify the ortholog with the highest degree of similarity.

The workflow was tested for its functionality and efficiency by integrating scRNA-seq datasets from heart failure patients with the corresponding mouse model. We were able to assign unique human orthologs to up to 80% of the mouse genes, utilizing the known 17,492 orthologous pairs. Curiously, the integration process enabled the identification of both common and unique regulatory pathways between species in heart failure.

In conclusion, our pipeline streamlines the integration process, enhances gene nomenclature alignment and simplifies the translation of mouse models to human disease. We have made the OrthoIntegrate R-package accessible on GitHub (https://github.com/MarianoRuzJurado/OrthoIntegrate), which includes the assignment of ortholog definitions for human and mouse, as well as the pipeline for integrating single cells.

**Keypoints:**

- Novel integration workflow for scRNA-seq data from different species in an easy to use R-package (“OrthoIntegrate”).
- Improved one-to-one ortholog assignment via sequence similarity scores and string similarity calculations.
- Validation of “OrthoIntegrate” results with a case study of snRNA-seq from human heart failure with reduced ejection fraction and its corresponding mouse model

## Introduction

In recent years, single cell transcriptomics (scRNA-seq) has become a transformative approach for cellular and molecular biology research. This technique allows for the simultaneous quantification of genes and related pathways in thousands of individual cells, enabling a greater understanding of cellular heterogeneity and gene regulation. By applying clustering algorithms, scRNA-seq enables the definition of cell types with high resolution and accuracy [1,2]. In general, these methods are used to gain insights into altered biological states and thus to identify differences between healthy and diseased phenotypes at a single cell level.

In order to analyze these scRNA-seq data bioinformatically, integration pipelines were developed with the aim of normalizing and scaling data and combining the respective individual cells into clusters with regard to their expression pattern [2,3]. Many methods are available [4] and several scRNA-seq pipelines were developed to integrate different datasets from the same species [2,5,6]. Other methods have been implemented to integrate single cell atlases from different species, e.g by aligning protein sequences via blast [7]. But there is to this date no standardized and easy way to integrate scRNA-seq data collected from different species, which also uniquely assigns orthologs between the species, especially for the highly used combination of a mouse model and respective human patient data. In the past, for each type of experimental design, methods were developed that were suitable to determine orthologs. Graph-based models were developed to compare microarray data between species and detect gene activity, but other methods specializing in meta-analyses of gene expression patterns were also common [8][9]. These methods may provide an excellent solution for microarray data, but are not suitable for embedding cross-species single cell data into one dimensional space as they only integrate on mutual expression of common orthologs. To generate shared embeddings for single cell datasets of different species, one to one relationships are required. Additionally, they don’t provide uniquely assigned orthologous lists for further analysis in other data. Over time, deep learning algorithms, which predict mRNA expression levels from whole-genome DNA sequencing data, related to trustworthy and robust orthologous lists have also grown in importance. These models are able to predict a gene’s On/Off state as well as which of two compared orthologs is more highly expressed [10]. However, for direct comparisons of available scRNA-SEQ datasets from different species, these methods are not applicable. The routine utilization of publicly accessible orthologous catalogs sourced from established databases, such as Ensembl [11], Orthologous Matrix (OMA) [12], and InParanoid [13], has become a ubiquitous undertaking for cross-species studies. While providing numerous lists, these databases do not focus on providing one-to-one gene assignments, which are necessary for transcription comparisons between species.

These limitations and the highly increasing demand for comparison of single cell data of various organisms prompted us to develop a R package called “OrthoIntegrate”. It features a pipeline for data integration and ortholog assignment, allowing for simple integration of data from mouse models and human patients. For the ortholog assignment process, we implemented an algorithm in the workflow that adjusts the different nomenclature between species before the integration takes place. For this purpose, we use the databases of Ensembl, NCBI, and Uniprot. [11,14,15]. The Ensembl database provides an excellent provisional assignment of orthologs, but there are, as previously mentioned, usually multiple assignments that cannot be used in the integration. To circumvent this problem, we additionally perform a nucleotide sequence alignment and a protein sequence alignment, using the Needleman-Wunsch global alignment algorithm [16] on sequences from the NCBI and Uniprot database.

Overall, we propose a bioinformatic pipeline to integrate single cell and nuclei sequencing data from mouse and humans to bioinformatically assess easily the full set of scRNA-seq tasks, like quality control (QC) steps, dimensionality reduction, clustering, cell-marker detection, differential gene expression and gene ontology analysis. We hope to simplify the analysis and the comparison of human patient data and mouse model data with this user-friendly pipeline and our ortholog finding algorithm.

To test the functionality and to illustrate the benefits of our pipeline, we integrated data sets from human heart failure patients with preserved ejection fraction (HFrEF) and the corresponding mouse model with a permanent ligation of the left anterior descending artery (LAD).

## Methods

### Single cell pre-processing

Single-cell RNA-seq results were processed by CellRanger (10x Genomics) version 6.1.1 software. The first step consisted of demultiplexing and processing raw base count files by the implemented *mkfastq* tool. The human raw reads were mapped to the reference genome hg38 (GRCh38-2020) using Cellranger count, whereas the mouse raw reads were mapped to the reference genome mm10 (GRCm38-2020). The secondary data analysis was initiated by using the Seurat 4.1.0 package in R. The data sets were first combined into a Seurat object and then subjected to a filtering process. Barcodes with too low (< 300) or too high number of genes (> 6000) were sorted out and not considered further in the data analysis. In addition, barcodes with too low (< 500) and too high read counts (> 15000) were also sorted out. To further ensure that no apoptotic cells or doublets were analyzed, we discarded barcodes with a high percentage of mitochondrial content (> 5%). The filtered gene counts were then logarithmized and normalized according to the tutorial for data analysis with Seurat. Baseline characteristics for the samples can be found in Supplement Table 1.

### Ortholog assignment and sample integration

In order to ensure the integration of the individual single cell datasets, a function was written to assign mouse orthologs to the human nomenclature using gene transfer format (GTF) files provided by 10x Genomics (GRCh38 for human, GRCm38 for mouse). We wanted to detect only well annotated genes between species, therefore we filtered predicted genes out of our GTFs. The function consists of a total of five approaches, which resulted in a clear assignment. First, orthologs were determined using the R package biomaRt. This package enables the retrieval of Gene ID symbols for mouse and human genes stored in the Ensembl database. This assigned the majority of genes in our human GTF file to at least one ortholog. If there were several entries of possible orthologs in the Ensembl database, a protein sequence comparison was initiated, which obtained protein sequences from the Uniprot database for the human gene and the possible mouse orthologs. These sequences were then aligned using the R package Biostrings 2.60.2. The alignment score was calculated based on the Needleman-Wunsch global alignment algorithm and substitution matrices for nucleotide sequences or protein sequences to determine the correct ortholog in terms of amino acid sequence or nucleotide sequence identity. However, there were still Gene ID symbols which could not be uniquely assigned even using the Uniprot database. In order to be able to assign these as well, a further comparison with usage of the nucleotide sequence is initiated. For this purpose, the sequences for the human gene and for the possible mouse orthologs were obtained from the NCBI database and aligned analogously to the previous step and assigned to an ortholog depending on the highest alignment score. If these alignment steps are not successful, a score was calculated by comparing the human ID symbol and the mouse ID symbols of possible orthologs. For this the Levenshtein distance measurement algorithm was used. All orthologs found are compared with the gene ID symbols in the GTF files used for the mapping of human reads and mouse reads. These GTF files are also used for the last step regarding the ortholog assignment. It was assumed here that gene ID symbols for mouse genes are often annotated in the same way as for human genes, with the only difference being that there is only one capital letter at the beginning of the name. If the gene name determined in this way now occurs in the mouse GTF, it was set as an ortholog. With this globally applicable list of orthologs between species, the datasets were now filtered by these and then merged into one object using Seurat’s canonical correlation analysis (CCA) integration.

### Clustering, silhouette coefficient and annotation

To classify cells into clusters based on their expressed genes, we used the *FindNeighbors* and *FindClusters* (resolution parameter = 0.3) function implemented in Seurat. These clusters are determined by applying the shared nearest neighbors (SNN) clustering algorithm. Using the Uniform Manifold Approximation and Projection (UMAP) dimension reduction, we were able to visualize our calculated cell clusters and the species overlap between our samples.

Calculations of the silhouette coefficient are based on computing a distance matrix based on the cell embeddings matrix for principal component analysis (PCA) performed by Seurat. This distance matrix includes the information of cell-cell distance, which is necessary for calculating the silhouette coefficient with our calculated clusters in the function *silhouette* of the cluster package (version 2.1.4). Additionally, the coefficients of the samples were averaged for each object. The orthologous lists for OMA, Biomart and InParanoid were created by following their introductions on their tool descriptions and by using the same GTF files as before (GRCh38, GRCm38).

For the assignment of cell clusters to cell types, we used a reference object that we had previously manually annotated with marker genes from Tombor et al. 2021 [17]. Here, the R package SingleR can be used to adopt marker genes that were used for the previous annotation of clusters of the reference object. These are then transferred and compared to marker genes of the cell clusters of our object to be annotated. Thus, a reproducible annotation can be guaranteed with the help of an exactly annotated data set.

### Differential gene expression analysis and gene ontology analysis

Detection of differentially regulated genes (DEG) for the cell type specific clusters was performed by the hurdle model of the MAST package (version 1.20.0). Results were filtered by their Bonferroni-adjusted p-value (p.adj < 0.05). The totality of DEGs were represented by Sankey plots created with the R package networkD3 (version 0.4). DEGs were divided according to their species and cell type assignment and then visualized for DEGs with a positive Log2FC and separately in another plot, for DEGs with a negative Log2FC. Here, DEGs occurring in both human and mouse for the respective cell type have been pooled. Visualization was done in the form of a Circos plot (R package circlize 0.4.14).

Gene Set Enrichment Analysis (GSEA) was performed using the R package clusterProfiler (version 4.2.2). GSEA terms were calculated separately for each cell type. The terms were sorted according to the Benjamini-Hochberg adjusted p.value and evaluated according to their “enrichment distribution”, which gives information about the regulation of the genes in the described pathway. The GSEA results were plotted in ridge plots. Additionally, for genes described in the pathway, the standard error of the mean (SEM) bar plot was created (for their averaged UMIs) by using the R package ggplot2.

## Results

### Generation of unique one to one ortholog assignments

In order to determine the appropriate ortholog, we utilized the Needleman-Wunsch algorithm to perform a pairwise global alignment. This calculation determines alignment scores based on differences in the amino acid or nucleotide sequences. In case no orthologs were found, or neither a protein- or nucleotide sequence is available for a certain gene, a lowercase matching of the human gene is searched in the mouse gene database (Fig. 1A). The Ensembl database assigned a total of 21,428 mouse orthologs to our human gene ID symbols. While the biomaRt package could assign 77% (16,573) of them uniquely, our algorithm increased the number of assignments to 82% (17,504). In our analysis, we assigned 714 genes through protein sequence alignment and 89 through nucleotide sequence comparison. Furthermore, we identified 42 orthologous pairs using the Levenshtein distance and an additional 86 using our lowercase matching approach. We then proceeded by filtering our human and mice data by these orthologs in our pipeline and replaced the mice nomenclature by the human nomenclature for the corresponding samples (Fig. 1B). At the end, we assigned ∼80% of the mice genes to human orthologs (Supplement Table 2). The replacement of mouse gene names with the human ortholog allowed us to integrate the human patient data with the mouse model data into one Seurat object.

**Fig.1:**
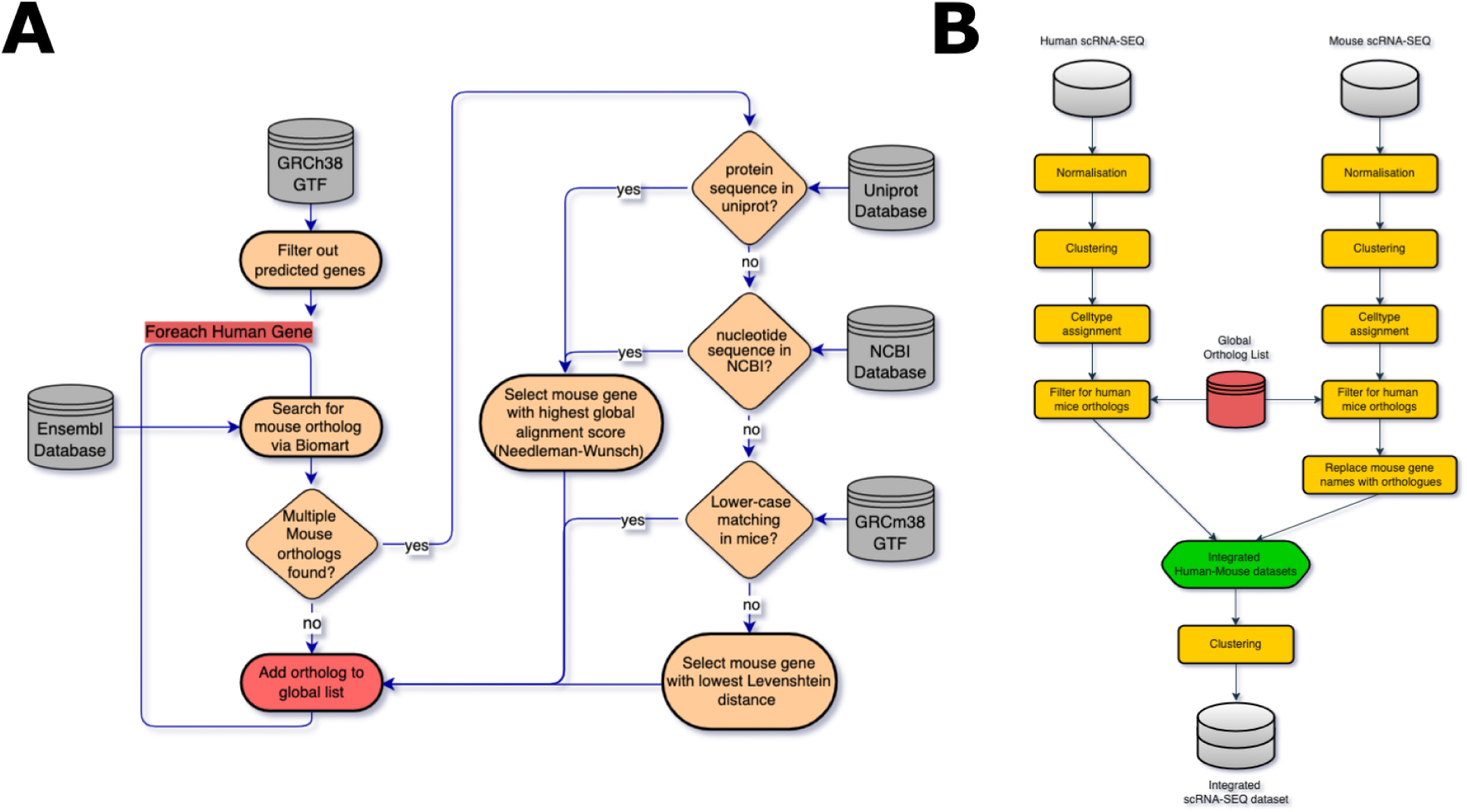
Integration process of human/mouse snRNA-SEQ data. (A) UML-Activity-Flowchart showing ortholog assignment pipeline for human to mouse gene symbols. First, the Gene transfer format file (GTF) for humans (GRCh38) is used to get all annotated gene nomenclatures. Then all genes are filtered out which are only predicted and not clearly detected. This list is now searched for orthologs using the Ensembl database; all 1:1 assignments can be included in our orthologous list. In the case of multiple assignments, all possible replacements are examined according to their protein sequence and an alignment score is calculated according to the global sequence alignment. If there is no protein sequence in the Uniprot database, the alignment score is calculated based on the nucleotide sequence using the NCBI database. Now the gene with the best result is set as an ortholog. All unassigned genes are additionally compared with the GTF file of GRCm38 using a lowercase matching and if there is a match, they will be added to the ortholog list. If all these approaches for a gene do not result in an ortholog, a Levenshtein distance score is calculated based on their gene names. (B) Single cell integration pipeline showing steps performed to integrate human and mouse scRNA-SEQ data in a joined UMAP projection. The scRNA-SEQ data from our human and mouse samples are first converted into Seurat objects and normalized. After that, clustering takes place and cell types can be determined. Using the orthologous list from our ortholog assignment algorithm, the objects can be subsetted according to the genes found and their nomenclature unified. This is followed by an integration into a single object and a clustering step.

### Comparison to other integration methods

We carefully inspected our data to determine species specific distribution by creating UMAP plots of all cells in our integrated object. Figure 2A shows that cells of mouse and human origin commingled in all clusters demonstrating the integration of the data sets. To find out how well our pipeline performs in integrating and clustering data based on our orthologous list, we compared our data object to other databases and tools and their orthologous pairs. For this purpose, we created Seurat objects using the same procedure as before with lists for orthologs from the bioinformatic tools: OMA, Biomart and InParanoid and visualized the species specific distribution of the cells in UMAP plots (Supplement Fig. 2). In order to assess the quality of the clustering, we calculated and visualized the mean value of the silhouette coefficient in a box plot (Fig. 2B, Supplement Table 3). The silhouette coefficient is particularly suitable because it gives a measure of quality of the clustering independent of the number of clusters. We can observe that our pipeline (OrthoIntegrate) has created the best integration achieving the highest silhouette coefficient and thus significantly improving the clustering compared to the other methods. Additionally, it is noteworthy that our pipeline achieved by far the most 1:1 protein coding and lncRNA coding orthologous pairs in comparison to the other described methods (Fig. 2C).

**Fig.2:**
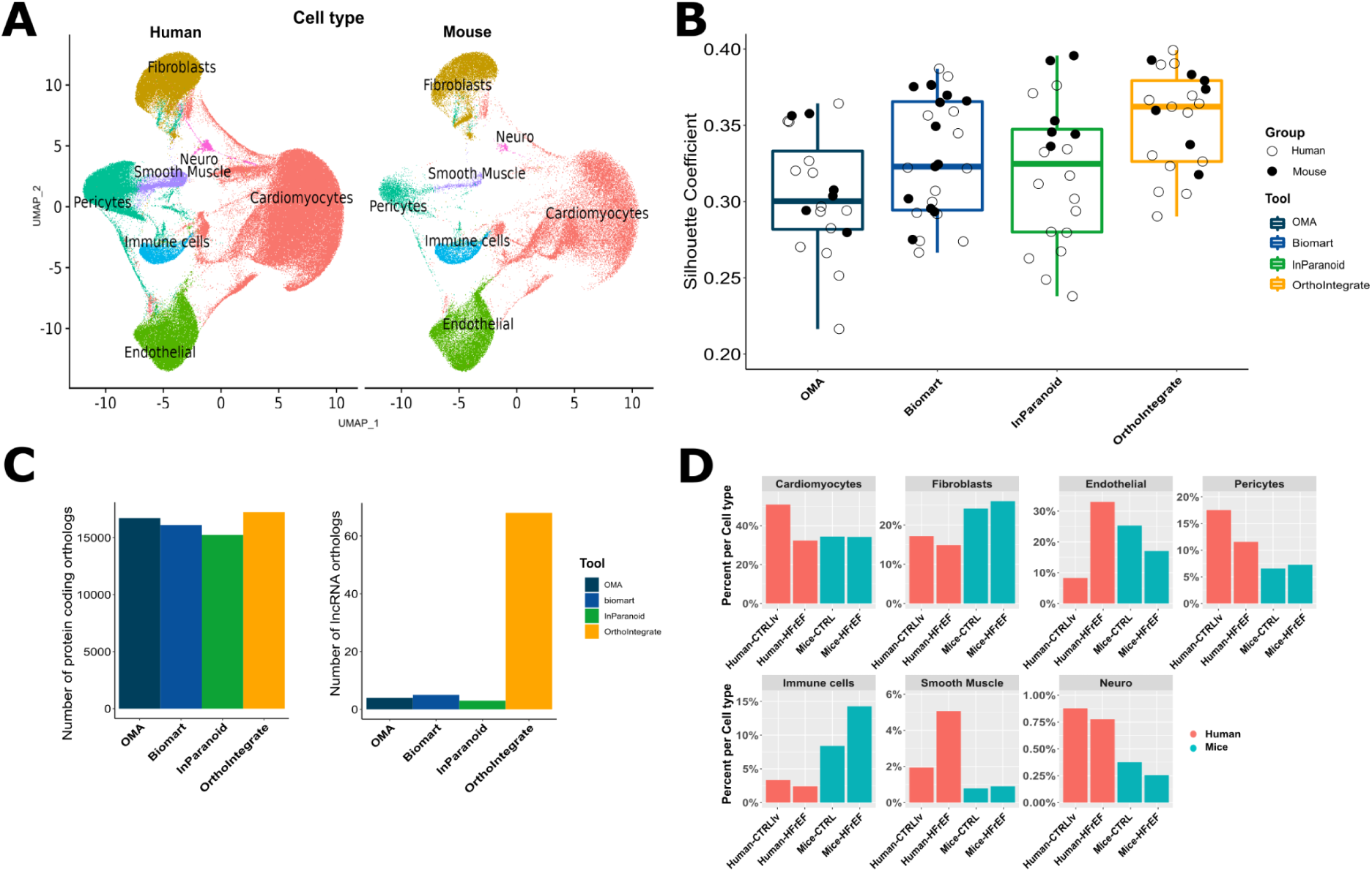
Integrated human and mouse scRNA-SEQ data of healthy and heart failure samples. (A) UMAP with defined clusters according to Seurat’s clustering, divided by species. Cells of mouse and human origin commingled in all clusters. There are no clusters formed that originated from only one of the two species. The cells were identified as cardiomyocytes (red), fibroblasts (yellow), endothelial cells (green), pericytes (turquoise), immune cells (blue), smooth muscle cells (purple) and neuronal cells (pink). (B) Box plot showing the average silhouette coefficient for clusterings based on different databases and tools. The dark blue box stands for the silhouette coefficient of the clustering made with an orthologous list using the tool OMA (Orthologous matrix). It is followed by the results for biomaRt (light blue), InParanoid (green) and our pipeline OrthoIntegrate (yellow). On the y-axis you can see the value of the Silhouette Coefficient. Additionally, each Silhouette Coefficient was calculated for each sample and depicted as a circle in their species specific color. (C) Bar plot with number of orthologs found which codes for a protein (left) and bar plot with number of orthologs found which codes for lncRNA. On the x-axis the used tool is depicted. (D) Bar plot showing cell composition of cell types in human (red) and mice (blue) samples. Samples were grouped based on their origin into human controls from the left ventricle (Human-CTRLlv), human HFrEF (Human-HFrEF), mouse controls (Mice-CTRL), and mouse HFrEF model (Mice-HFrEF). Cell types were then analyzed for their composition from the previously mentioned groups and plotted. The percent composition of the cell types are then shown as bar plots.

### Integration of patient data with mouse model data

In the integrated object no human or mouse specific clusters were identified, which indicates the common clustering of the different cell types. All clusters could be annotated by using specific marker genes and the R package SingleR into cardiomyocytes (CMs), pericytes (PCs), smooth muscle cells (SMCs), fibroblasts (FBs), endothelial cells (ECs), immune cells (ICs) as well as neuronal cells (NCs) (Fig. 2A). Of note, characteristic marker expression was similar for mice (Supplement Fig. 1A) and human cells (Supplement Fig. 1B). In addition, we analyzed how the distribution of cell types was affected by the biological state of the samples. We found that the number of human CMs decreases when we compare the control samples with the HFrEF samples (45% -> 25%). However, in mice, there is no difference in the number of CMs between model and healthy individuals (∼25%) (Fig. 2D). Furthermore, we see a marked difference in the distribution of ECs in our human patients compared with our mouse models. Here, we observed a significant increase in the EC population between healthy hearts from human patients (∼8%) and those with HFrEF (∼30%). In contrast, we noticed a decrease in ECs in mice upon HFrEF (from 25% in controls to 18% in HFrEF). In addition, minor changes are also observed in the numbers of other cell types.

### Differential gene expression between mice and man in up-regulated genes

Next, we compared the regulation of gene expression by HFrEF in mice and humans. The results of the differentially expressed gene (DEG) analysis, showed that there were clear similarities and differences regarding the expression of genes in general and in the respective cell types between species. In the human samples 4,141 genes were differentially expressed between HFrEF and the control condition. From these, 2,995 were up-regulated and 1,146 down-regulated. In the mouse data, 4,654 genes were significantly different, of which 3,699 were up-regulated and 955 down-regulated (Supplement Table 4).

A cell type- and species-specific DEG analysis allowed us to identify cell type-specific expression patterns between human patients and mouse models (Fig. 3). First, we analyzed the extent to which the up-regulated DEGs are specific to each species or which DEGs are found in both species and the distribution in each cell type (Figs. 3A, 3B). When analyzing the cardiomyocyte (CM) subpopulation, we found 2,906 DEGs are up-regulated in human CMs, of which 62% (1,806) were also significantly up-regulated in mice CMs. The other 38% (1,100) of the DEGs were uniquely upregulated in the human HFrEF versus human control samples. In the HFrEF mouse model, we found 3,908 DEGs which were not up-regulated in the human patient samples. Observation of the downregulated CM genes showed that 1,083 genes are significantly down-regulated upon HFrEF. From these 82% (888) were exclusively regulated in humans. The remaining 195 genes (18%) were also significantly downregulated in the mouse models. In addition, we observed that 593 DEGs are specific for the HFrEF mouse model.

**Fig.3:**
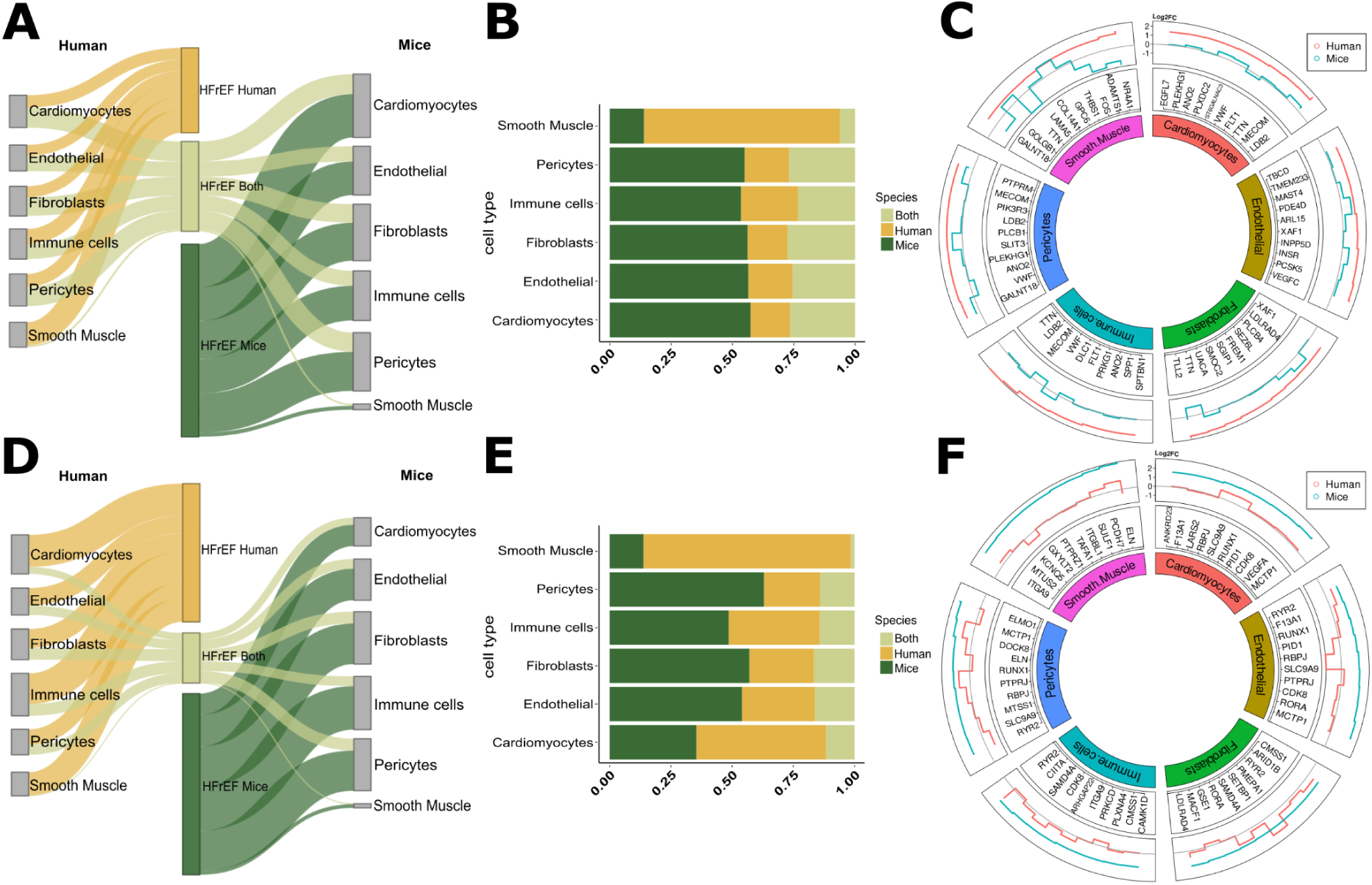
DEG analysis shows similar and different expressed DEGs. (A) Sankey plot illustrating the distribution of up-regulated differentially regulated genes (DEG) in the corresponding cell types. The width of the paths illustrates the number of DEGs that are either human specific (yellow) are detected in both species (light green) or are mouse-specific (dark green). DEG analysis was performed for each cell type individually. It should be noted that neuronal cells were omitted from all further analyses due to their insufficient number of cells in the mouse data. (B) Bar plot showing the percentage distribution of up-regulated genes in the respective cell types. The y-axis represents the cell type, while the x-axis displays the percentage distribution (1.00 = 100%). The bars are stacked on top of each other and color coded for easy interpretation (yellow = human specific, light green = both species, dark green = mouse specific). (C) Circos plot showing the ten most up-regulated genes for the comparison between human HFrEF and the corresponding control, seperated for all cell types. The outer ring of the plot shows the values for the changes in gene expression (Log2FC) for the human (red line) in comparison to the regulation in mouse HFrEF samples (blue line). In the middle ring, the corresponding gene name is listed. In the inner ring, the cell types are shown. (D) Sankey plot similar to (A) representing down-regulated DEGs. (E) Stacked bar plot similar to (B) showing the percentage distribution of down-regulated genes in the respective cell types. (F) Circos plot similar to (C) illustrating the ten most down-regulated genes in mice HFrEF samples in comparison to the regulation of these genes in humans.

In ECs, we observed 2,339 up-regulated genes in our human HFrEF, of which 42% (971) of the genes were specific for our cardiac patients. In the mouse model 4,399 DEGs were detected. From these 3,031 (69%) DEGs were exclusively regulated in the mice samples. With respect to the down-regulated DEGs, we observed that 749 genes are downregulated in our human HFrEF, of which 44% (485) of the DEGs are only found in human patient data. The remaining 264 genes were also present in the mice DEGs. Additionally, we found 880 genes which were only regulated in mouse HFrEF models.

Notably we found far less DEGs in the mouse SMCs in contrast to the human samples. However, this could be related to the total number of SMCs in mice, which is far less compared to the human samples (Figs. 3A, 3C). Looking at the distribution of DEGs in all cell types, we can see that the percentage of commonly regulated genes is smaller when comparing the down-regulated DEGs to the upregulated DEGs (Figs. 3B, 3E).

Figure 3C and Figure 3F show the highest upregulated genes per cell type in humans and mice along with the regulation of that gene in the other species. Hereby, we can observe how the genes with the largest changes in human heart failure patients behave in the respective mouse model.

We observed that the expression of the most regulated genes in human cell types is mostly much less affected in the mouse models. For example, we see a gene of the LIM-Domain family, *LDB2*, in human CMs as the gene with the most change in expression (Log2FC = 2.15). These genes are well known as adapter molecules which allow assembly of transcriptional regulatory complexes in CM. In contrast, only minimal up-regulation of the gene is noticeable in mouse HFrEF models (Log2FC = 0.38). Other genes such as the VEGF receptor *FLT1*, which is well detected in cardiomyocytes in human cardiac tissue, show a negative log2FC (expression values decreased in comparison to the corresponding control) in mice CMs but increased expression in human CM, so these genes are regulated in a different direction than in human patients. However, there are also genes that share similar regulation in their respective cell types. Thus, we can observe that Phosphodiesterase 4D (*PDE4D)* and ADP Ribosylation Factor Like GTPase 15 (*ARL15)* show a similar change in ECs with respect to their expression as in human ECs. In the ten most upregulated genes in the mouse model data, we can observe three genes that also show a significant increase in their expression in humans (*RBPJ, SLC9A9, RUNX1*). The other genes, however, show little to no change. If we now observe the expression changes in ECs, there are DEGs showing an opposite direction in their expression change (*RBPJ, PID1, SLC9A9*). These different and similar gene expressions in the cell types are first indicators of differences and similarities between human patients and mouse models.

### Pathway enrichment results

Since we found an unexpectedly high number of differentially regulated genes, we investigated if this might indicate overall changes in pathways and pathological processes or whether the difference relates more to the alternative use of genes with similar functions in mice and humans. Therefore, we investigated how the enriched signaling pathways differ in cardiomyocytes. We observed that signaling pathways mainly dealing with energy metabolism are similarly and significantly down-regulated in patients with heart disease as well as in mouse models (Fig. 4A). The genes included in these pathways, such as “ATP metabolic process”, “mitochondrial respiratory chain complex I assembly”, “electron transport chain” and “cellular respiration”, show significant down-regulation compared to their corresponding control (Fig. 4B). These data suggest a conservation of disturbed mitochondrial metabolism in both mice and humans in heart failure. In contrast, larger differences are observed for the up-regulated gene sets. Among the most regulated pathways specifically detected in the human data set, we noted the terms “angiogenesis” and “regulation of small GTPase mediated signal transduction” (Fig. 4C). These gene sets are not found among the regulated pathways in mice (Supplement Table 5). Examples for angiogenesis-related genes, which are specifically induced in human heart failure but not in mice models, include receptors such as the *VEGF*-receptor FLT1, or transcription factors like the mesenchyme homeobox protein 2 (MEOX2) (Figure 4D). In addition, many GTPase regulatory genes were found specifically increased in humans, including *MCF2L* and *RASGRF2*, which are known to regulate *RAC1*, and *SPATA13*, which enables guanyl-nucleotide exchange factor activity.

**Fig.4:**
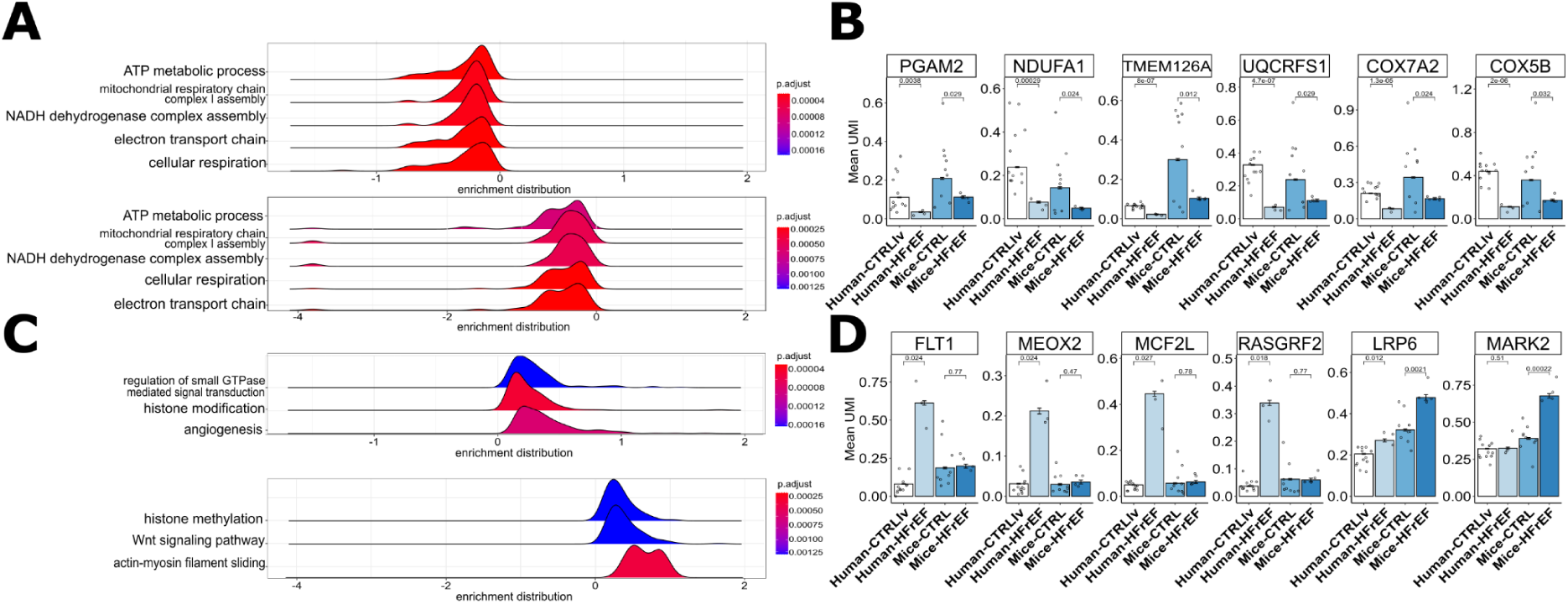
GSEA analysis shows regulated pathways upon heart failure in human and mouse cardiomyocytes. (A) Ridge plot visualizing five of the same down-regulated pathways found in human (top plot) and mouse (underneath plot) cardiomyocytes, while HFrEF is present. The y-axis displays the description of the identified term, while the x-axis shows the enrichment distribution, which indicates the pathway’s overall regulation (enrichment distribution > 0 implies up-regulation, and < 0 indicates down-regulation). The ridges are also color-coded indicating significance of the found term (B) Bar plot with mean values for the amount of unique molecular identifiers (UMIs) in the cells for the depicted genes. The genes are found to be down-regulated in mice and humans in the pathways shown in (A). On the x-axis, the samples are grouped for their biological condition (CTRLlv = Control left ventricle) and species. The values for the individual samples are depicted as circles. Additionally, t-test values are shown over the bars for the tested group. (C) Ridge plot visualizing similar to (A) three of the different up-regulated pathways found for human cardiomyocytes and cice cardiomyocytes. On the y-axis there is the description of the found term and on the x-axis is the enrichment distribution depicted. (D) Bar plot similar to (A) with mean values for UMIs in the cells for the depicted genes found in terms of (C).

On the other hand, our analysis revealed that pathways such as “Wnt signaling pathway” and “actin-myosin filament sliding” are up-regulated specifically in the mouse HFrEF model (Fig. 4C). Genes associated with Wnt signaling include *LRP6*, a known inhibitor of cardiomyocyte proliferation, and the serine/threonine-protein kinase *MARK2*, which regulates the stability of microtubules through phosphorylation and inactivation of several microtubule-associated proteins [18].

## Discussion

The ever growing number of published single cell experiments enables scientists to deepen our knowledge about transcriptional changes of individual cell types as well as species specific regulatory changes, upon disease conditions. Particular combination of single cell datasets from different species in the same UMAP projection allows the detection of well conserved or species specific regulatory networks [19–21]. However, to integrate datasets from different species a well curated list of orthologous and related genes has significant advantages.

Here we propose the R-package OrthoIntegrate that enables scientists to integrate single cell datasets from different species into a shared dimensional space. To generate high quality and uniquely mapped orthologous lists between different species, we implemented an improved orthologous assignment pipeline which results in up to 10% more uniquely assigned orthologs between human and mouse in contrast to the Ensembl orthologous list (Biomart). Furthermore, the package contains functions which use our advanced orthologous assignments and easily combines Seurat objects from humans and mice. Moreover, it is highly adaptable and can be easily customized to support other species as well.

We demonstrated the usability of combining cross-species single cell datasets on a heart failure dataset with reduced ejection fraction of humans and mice. Hereby, we could show that there are major differences in the cell type expression patterns, which are differentially regulated in humans or mice upon HFrEF. Yet there are also commonly regulated pathways that reflect an evolutionary conserved transcriptomic answer to severe damages in heart cells. These include the down-regulation of important mitochondrial metabolic pathways, which provide ATP for the heart, which are down-regulated in mice and human heart failure. The adult heart is the most energy consuming organ, and a critical role of mitochondrial function in maintaining a healthy heart is well known [22].

However, there were also interesting differences between human and mice heart failure samples. Surprisingly, in humans, but not in mice, many genes associated with “angiogenesis” were induced in cardiomyocytes. For example, the VEGF receptor FLT1 was significantly induced in the human samples. FLT1 is well known to primarily mediate VEGF signaling in endothelial cells, but its role in cardiomyocytes is less clear [23]. FLT1 protein is well detected in cardiomyocytes in human cardiac tissue [24]. Functionally, FLT1 was shown to partially mediate VEGF-induced cardiomyocyte differentiation of embryonic stem cells [25] and mediates VEGF induced cardiomyocyte calcium signaling and contractility in the embryonic zebrafish heart [26]. Cardiomyocyte specific deletion of Flt-1 was shown to worsen cardiac remodeling and hypertrophy induced by pressure overload [27], suggesting that the up-regulation of its expression in humans may represent a compensatory cardioprotective mechanism. A second example is MEOX2, also known as GAX, which was assigned the GO term Angiogenesis because of its role in endothelial fatty acid transport [28]. MEOX2 plays a critical role in development of all muscle lineages [29]. In cardiomyocytes, MEOX2 overexpression blocks proliferation during heart morphogenesis causing proliferating cardiomyocytes to withdraw from the cell cycle [30]. In addition, various guanine nucleotide exchange factors and regulators of Rho/Rac signaling pathways were shown to be specifically induced in human cardiomyocytes. While G protein-coupled signaling is well known to control cardiomyocytes [31], the function of the highly induced regulatory genes identified here (e.g. MCF2L, RASGRF2, and SPAT13) have not been studied in cardiomyocytes.

In mice, a predominant expression of genes associated with Wnt signaling were detected. Although the majority of the identified genes has not been directly linked to cardiomyocyte-specific functions, Wnt signaling is critically regulating cardiac hypertrophy, remodeling and regeneration [32]. Therefore, these findings as well as the other identified species-specific pathways deserve a closer in depth validation and investigation.

## Limitations

The main limitation of our ortholog assignment and sample integration pipeline, however, is the dependence on reliable databases for orthologous lists. Another problem with this approach is that it fails to take biological functions of the possible orthologs into account and decide based on the most suitable function which ortholog may fit the best and not only on sequence similarity.

## Acknowledgement

The study was supported by grants from the German Centre for Cardiovascular Research (DZHK) to D.J. and S.D., the German Research Foundation (DFG; Exc2026/1) and the Dr. Rolf M. Schwiete Stiftung, Projekt 08/2018 to S.D.

## Code availability

The OrthoIntegrate package containing the integration pipeline and the ortholog algorithm are available on Github (github.com/MarianoRuzJurado/OrthoIntegrate). Additionally, codes for R analysis and plots of data presented in this study are available on another GitHub repository (github.com/MarianoRuzJurado/BriefinBio_RuzJurado_et_al_2023).

## Ethics declaration

The authors declare no competing interests.

## Supplementary Figures

**Supplement Fig. 1:**
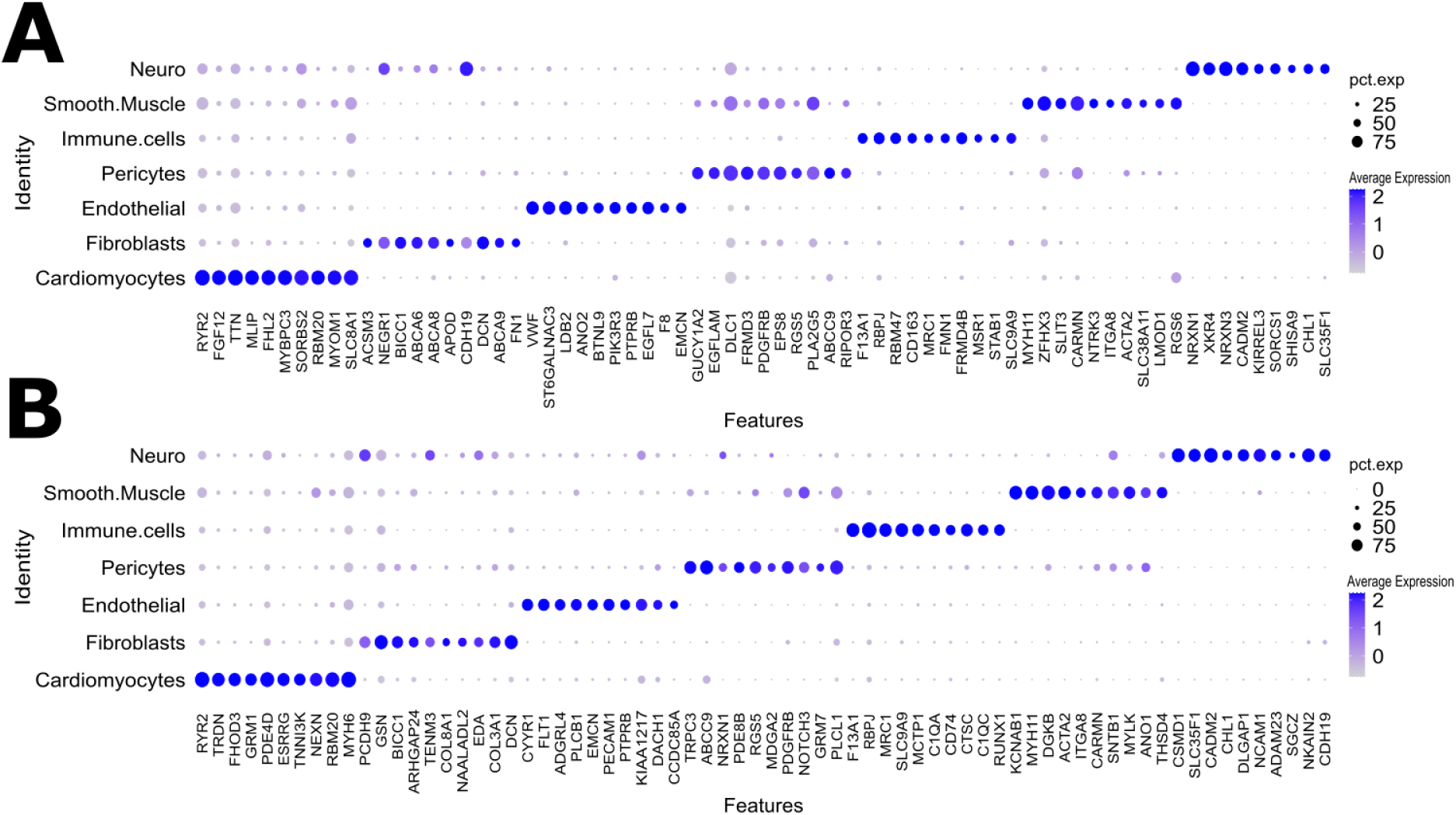
Known cell type marker genes are found for human and mice data. (A) Dot plot depicting the average expression levels and expression proportions in human samples of the top ten feature genes for the found cell types. The size of the dot represents the proportion of cells expressing the indicated gene within a cell type, and the color indicates the average expression level of cells. (B) Dot plot depicting the average expression levels and expression proportions in mice samples of the top ten feature genes for the found cell types. Similar to (A) the size of the dot represents the proportion of cells expressing the indicated gene within a cell type, and the color indicates the average expression level of cells.

**Supplement Fig. 2:**
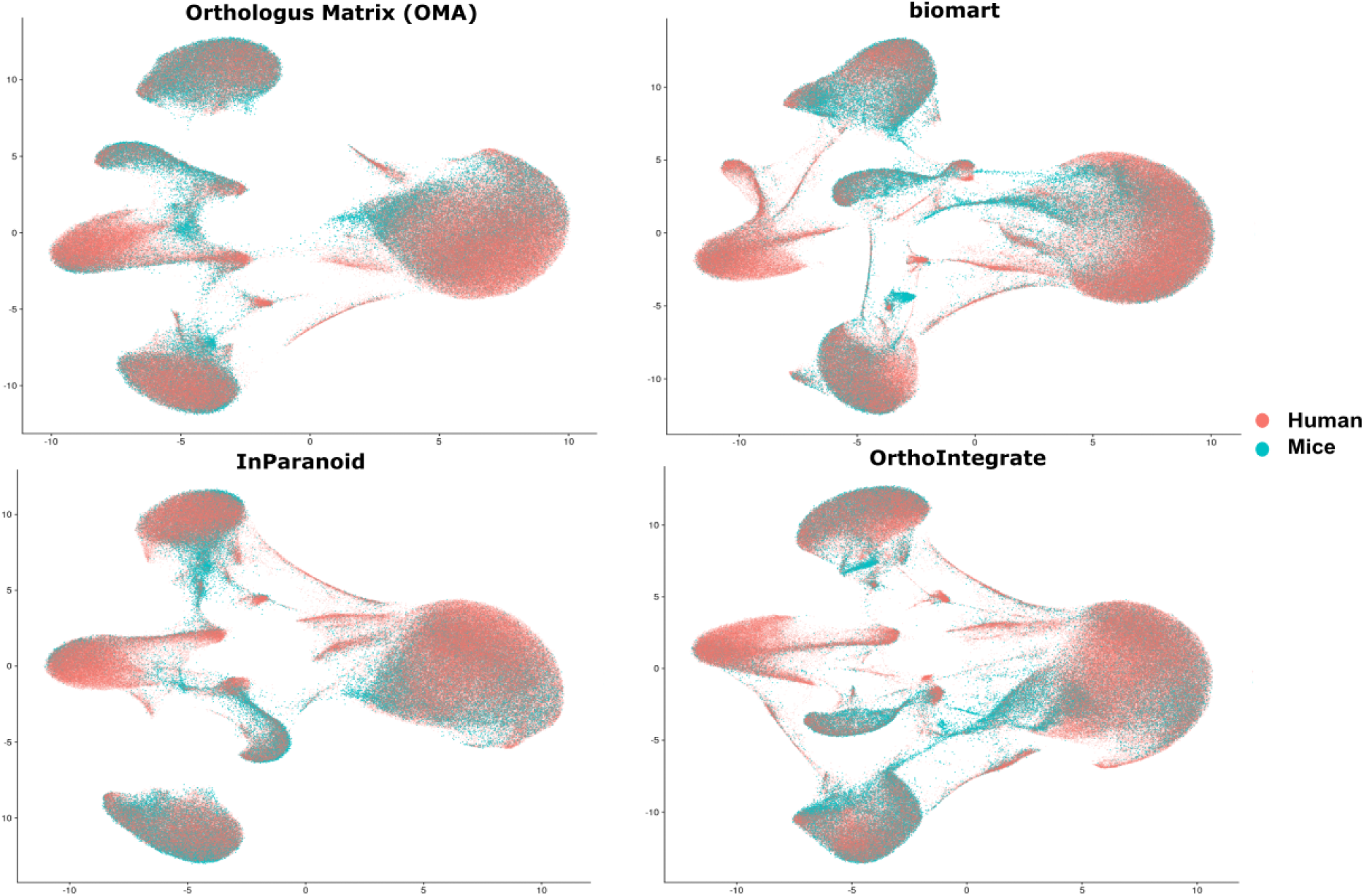
Overlapping of human and mice cells after Seurat integration with tool specific orthologous list. UMAPs showing human cells (red) and mice cells (blue) in a common UMAP projection for each tool used for integrating the data. First UMAP was performed on an object made with an orthologous list of OMA, followed by Biomart and InParanoid. The last UMAP shows the projection for the OrthoIntegrate pipeline.

## References

1. Jovic D, Liang X, Zeng H, Lin L, Xu F, Luo Y. Single-cell RNA sequencing technologies and applications: A brief overview. Clin Transl Med. 12:e6942022;

2. Stuart T, Butler A, Hoffman P, Hafemeister C, Papalexi E, Mauck WM 3rd, et al. Comprehensive Integration of Single-Cell Data. Cell. 177:1888–902.e212019;

3. Hwang B, Lee JH, Bang D. Single-cell RNA sequencing technologies and bioinformatics pipelines. Exp Mol Med. 50:1–142018;

4. Ziegenhain C, Vieth B, Parekh S, Reinius B, Guillaumet-Adkins A, Smets M, et al. Comparative Analysis of Single-Cell RNA Sequencing Methods. Mol Cell. 65:631–43.e42017;

5. Parekh S, Ziegenhain C, Vieth B, Enard W, Hellmann I. zUMIs - A fast and flexible pipeline to process RNA sequencing data with UMIs. Gigascience. 2018; doi: 10.1093/gigascience/giy059.

6. Korsunsky I, Millard N, Fan J, Slowikowski K, Zhang F, Wei K, et al. Fast, sensitive and accurate integration of single-cell data with Harmony. Nat Methods. Nature Publishing Group; 16:1289–962019;

7. Tarashansky AJ, Musser JM, Khariton M, Li P, Arendt D, Quake SR, et al. Mapping single-cell atlases throughout Metazoa unravels cell type evolution. Elife. 2021; doi: 10.7554/eLife.66747.

8. Lu Y, Rosenfeld R, Nau GJ, Bar-Joseph Z. Cross Species Expression Analysis of Innate Immune Response. In: Batzoglou S, editor. Research in Computational Molecular Biology. Berlin, Heidelberg: Springer Berlin Heidelberg; p. 90–107.

9. Kristiansson E, Österlund T, Gunnarsson L, Arne G, Larsson DGJ, Nerman O. A novel method for cross-species gene expression analysis. BMC Bioinformatics. 14:702013;

10. Washburn JD, Mejia-Guerra MK, Ramstein G, Kremling KA, Valluru R, Buckler ES, et al. Evolutionarily informed deep learning methods for predicting relative transcript abundance from DNA sequence. Proc Natl Acad Sci U S A. 116:5542–92019;

11. Cunningham F, Allen JE, Allen J, Alvarez-Jarreta J, Amode MR, Armean IM, et al. Ensembl 2022. Nucleic Acids Res. 50:D988–952022;

12. Altenhoff AM, Train C-M, Gilbert KJ, Mediratta I, Mendes de Farias T, Moi D, et al. OMA orthology in 2021:p website overhaul, conserved isoforms, ancestral gene order and more. Nucleic Acids Res. 49:D373–92021;

13. Persson E, Sonnhammer ELL. InParanoid-DIAMOND: faster orthology analysis with the InParanoid algorithm. Bioinformatics. 38:2918–92022;

14. Sayers EW, Bolton EE, Brister JR, Canese K, Chan J, Comeau DC, et al. Database resources of the national center for biotechnology information. Nucleic Acids Res. 50:D20–62022;

15. UniProt Consortium. UniProt: the universal protein knowledgebase in 2021. Nucleic Acids Res. 49:D480–92021;

16. Needleman SB, Wunsch CD. A general method applicable to the search for similarities in the amino acid sequence of two proteins. J Mol Biol. 48:443–531970;

17. Tombor LS, John D, Glaser SF, Luxán G, Forte E, Furtado M, et al. Single cell sequencing reveals endothelial plasticity with transient mesenchymal activation after myocardial infarction. Nat Commun. 12:6812021;

18. Wu Y, Zhou L, Liu H, Duan R, Zhou H, Zhang F, et al. LRP6 downregulation promotes cardiomyocyte proliferation and heart regeneration. Cell Res. 31:450–622021;

19. Balachandran S, Pozojevic J, Sreenivasan VKA, Spielmann M. Comparative single-cell analysis of the adult heart and coronary vasculature. Mamm Genome. 2022; doi: 10.1007/s00335-022-09968-7.

20. Butler A, Hoffman P, Smibert P, Papalexi E, Satija R. Integrating single-cell transcriptomic data across different conditions, technologies, and species. Nat Biotechnol. 36:411–202018;

21. Baron M, Veres A, Wolock SL, Faust AL, Gaujoux R, Vetere A, et al. A Single-Cell Transcriptomic Map of the Human and Mouse Pancreas Reveals Inter- and Intra-cell Population Structure. Cell Syst. 3:346–60.e42016;

22. Huss JM, Kelly DP. Mitochondrial energy metabolism in heart failure: a question of balance. J Clin Invest. 115:547–552005;

23. Kurotsu S, Osakabe R, Isomi M, Tamura F, Sadahiro T, Muraoka N, et al. Distinct expression patterns of Flk1 and Flt1 in the coronary vascular system during development and after myocardial infarction. Biochem Biophys Res Commun. 495:884–912018;

24. Karlsson M, Zhang C, Méar L, Zhong W, Digre A, Katona B, et al. A single-cell type transcriptomics map of human tissues. Sci Adv. 2021; doi: 10.1126/sciadv.abh2169.

25. Chen Y, Amende I, Hampton TG, Yang Y, Ke Q, Min J-Y, et al. Vascular endothelial growth factor promotes cardiomyocyte differentiation of embryonic stem cells. Am J Physiol Heart Circ Physiol. 291:H1653–82006;

26. Rottbauer W, Just S, Wessels G, Trano N, Most P, Katus HA, et al. VEGF-PLCgamma1 pathway controls cardiac contractility in the embryonic heart. Genes Dev. 19:1624–342005;

27. Mei L, Huang Y, Lin J, Chu M, Hu C, Zhou N, et al. Increased cardiac remodeling in cardiac-specific Flt-1 receptor knockout mice with pressure overload. Cell Tissue Res. 362:389–982015;

28. Coppiello G, Collantes M, Sirerol-Piquer MS, Vandenwijngaert S, Schoors S, Swinnen M, et al. Meox2/Tcf15 heterodimers program the heart capillary endothelium for cardiac fatty acid uptake. Circulation. 131:815–262015;

29. Skopicki HA, Lyons GE, Schatteman G, Smith RC, Andrés V, Schirm S, et al. Embryonic expression of the Gax homeodomain protein in cardiac, smooth, and skeletal muscle. Circ Res. 80:452–621997;

30. Fisher SA, Siwik E, Branellec D, Walsh K, Watanabe M. Forced expression of the homeodomain protein Gax inhibits cardiomyocyte proliferation and perturbs heart morphogenesis. Development. 124:4405–131997;

31. Brown JH, Del Re DP, Sussman MA. The Rac and Rho hall of fame: a decade of hypertrophic signaling hits. Circ Res. 98:730–422006;

32. Bergmann MW. WNT signaling in adult cardiac hypertrophy and remodeling: lessons learned from cardiac development. Circ Res. 107:1198–2082010;

